# Severe corrosion of carbon steel in oil field produced water can be linked to methanogenic archaea containing a special type of [NiFe] hydrogenase

**DOI:** 10.1101/2020.07.23.219014

**Authors:** Sven Lahme, Jaspreet Mand, John Longwell, Ramsey Smith, Dennis Enning

**Author notes:** Address correspondence to Sven Lahme, and Dennis Enning.

## Abstract

Methanogenic archaea have long been implicated in microbially influenced corrosion (MIC) of oil and gas infrastructure, yet a first understanding of the underlying molecular mechanisms has only recently emerged. We surveyed pipeline-associated microbiomes from geographically distinct oil field facilities and found methanogens to account for 0.2 – 9.3% of the sequenced communities. Neither the type nor the abundance of the detected methanogens correlated to the perceived severity of MIC in these pipelines. Using fluids from one pipeline, MIC was reproduced in the laboratory, both under stagnant conditions and in customized corrosion reactors simulating pipeline flow. High corrosion rates (up to 2.43 mm Fe^0^ yr^−1^) with macroscopic, localized corrosion features were attributed to lithotrophic, mesophilic microbial activity. Other laboratory tests with the same waters yielded negligible corrosion rates (< 0.08 mm Fe^0^ yr^−1^). Recently a novel [NiFe] hydrogenase, from *Methanococcus maripaludis* strain OS7, was demonstrated to accelerate corrosion. We developed a specific qPCR assay and detected the gene encoding the large subunit of this hydrogenase (labelled *micH*) in corrosive (> 0.15 mm Fe^0^ yr^−1^) biofilms. The *micH* gene on the other hand was absent in non-corrosive biofilms despite an abundance of methanogens. Reconstruction of a nearly complete *Methanococcus maripaludis* genome from a highly corrosive mixed biofilm revealed *micH* and associated genes in near-identical genetic configuration as strain OS7, thereby supporting our hypothesis that the encoded molecular mechanism contributed to corrosion. Lastly, the proposed MIC biomarker was detected in multiple oil fields, indicating a geographically widespread involvement of this [NiFe] hydrogenase in MIC.

**IMPORTANCE:** Microorganisms can deteriorate built environments, which is particularly problematic in the case of pipelines transporting hydrocarbons to industrial end users. MIC is notoriously difficult to detect and monitor and as a consequence, is a particularly difficult corrosion mechanism to manage. Despite the advent of molecular tools and improved microbial monitoring strategies for oil and gas operations, specific underlying MIC mechanisms in pipelines remain largely enigmatic. Emerging mechanistic understanding of methanogenic MIC derived from pure culture work allowed us to develop a qPCR assay that distinguishes technically problematic from benign methanogens in a West African oil field. Detection of the same gene in geographically diverse samples from North America hints at the widespread applicability of this assay. The research presented here offers a step towards a mechanistic understanding of biocorrosion in oil fields and introduces a binary marker for (methanogenic) MIC that can find application in corrosion management programs in industrial settings.

## INTRODUCTION

Microbially influenced corrosion (MIC) is problematic to oil field operations when it leads to extensive damage requiring the repair or replacement of infrastructure such as carbon steel pipelines. While some metal loss is expected and accounted for during the design of the infrastructure (e.g. 0.1 mm yr^−1^ for a pipeline designed with 3 mm corrosion allowance for 30 years design life), higher rates of corrosion, unless detected and mitigated in a timely manner, can necessitate costly repairs or replacements long before the design service life has been actualized.

The most common oil field corrosion mechanisms are linked to CO_2_ and/or hydrogen sulfide (H_2_S) which are present in oil-bearing formations and readily dissolve into the water that is associated with production fluids (termed produced water). The deleterious effects of these acid gases are managed through injection of corrosion inhibitors (CIs), typically film-forming surfactants that act as a diffusion barrier between steel and the aqueous corrosive environment (1, 2). Acid gas corrosion can be modeled *in silico* (3) or simulated in laboratory reactors (4) to understand the rates of metal loss expected under specific field conditions (i.e. flow, temperature, brine chemistry and partial pressure of acid gases). Furthermore, laboratory testing provides a robust approach to ensure CI effectiveness (4, 5). Microbial corrosion, on the other hand, cannot be adequately modeled and laboratory qualification of oil field biocides, applied to control MIC, typically falls short at predicting product effectiveness in the field. As a consequence, field-based detection and monitoring is particularly important for effective management of this corrosion mechanism.

In environments devoid of oxygen, MIC has been linked to a variety of anaerobic microorganisms ((6) and references therein); yet most attention has historically been given to the sulfate-reducing bacteria (SRB) due to their well-documented ability to cause severe corrosion in laboratory studies and their prevalence in production systems ((7) and references therein). SRB impact metal integrity by producing hydrogen sulfide (H_2_S) as a corrosive metabolic end-product of sulfate reduction with (usually) organic electron donors. A small number of lithotrophic SRB isolates have further been shown to severely accelerate corrosion of steel in laboratory tests by utilizing cathodic electrons for their metabolism (8–10). It was proposed that outer membrane c-type cytochromes enable the direct uptake of electrons from steel in one such isolate, *Desulfovibrio ferrophilus* strain IS5 (11). In addition to SRB, also methanogenic archaea are increasingly viewed as prime culprits in the corrosion of steel (12–19). Some methanogens have been shown to utilize metallic iron (Fe^0^) in carbon steel as the sole electron donor for methanogenesis (13, 14, 19, 20). Electrochemical studies suggested a mediator-free direct electron uptake mechanism for one of these Fe^0^-utilizing isolates, *Methanobacterium-like* strain IM1 (21). The potential technical relevance of this microorganism has been demonstrated in laboratory studies where strain IM1 was grown in flow-through systems (22).

A deeper understanding of the underlying biochemical basis of Fe^0^ utilization by methanogenic archaea has recently been provided. By working with filtered spent culture medium as well as hydrogenase-negative mutants of *Methanococcus maripaludis* strain MM901, Deutzmann and colleagues demonstrated that free, steel-associated hydrogenases from this strain catalyzed the production of hydrogen (H_2_) from Fe^0^ granules (23). Whether these hydrogenases originated from partial lysis of the laboratory culture (e.g. in stationary phase) or were actively excreted is currently unknown. Likewise, it is unclear whether this mechanism can produce technically relevant MIC in laboratory or oil field settings. Tsurumaru and colleagues performed comparative genomics on multiple strains of *Methanococcus maripaludis* and identified a 12 kb genomic region (termed ‘MIC island’) that was unique to pure cultures of strain OS7, the only strain of *M. maripaludis* that accelerated Fe^0^ oxidation (corrosion) in their tests (24). This ‘MIC island’ encoded for a novel type of [NiFe] hydrogenase along with an extracellular transport system. It was postulated that the secreted [NiFe] hydrogenase would catalyze the reduction of H^+^ to molecular hydrogen on iron surfaces, thereby accelerating the oxidation of the provided iron granules.

These recent insights into the molecular mechanisms of MIC by methanogenic archaea prompted us to investigate the role of methanogens in the corrosion of oil and gas infrastructure. Since MIC is a biofilm-associated phenomenon (25), we began by surveying the archaeal microbiomes of steel-associated solids from pipelines in North America, East Asia and West Africa. We then performed a comprehensive study of MIC using produced waters from one of these locations in controlled laboratory reactors that allowed for the growth of methanogenic archaea. The extent of corrosion as well as associated microbial communities were investigated under various enrichment conditions with natural oil field waters, as well as synthetic brines to specifically enrich for lithotrophic microorganisms. Building on these laboratory tests, we gained insights into the molecular mechanisms of MIC through shotgun DNA sequencing and metagenomic analysis of a highly corrosive, lithotrophic biofilm grown under simulated pipeline conditions. Lastly, we developed a novel qPCR assay for the detection of the abovementioned [NiFe] hydrogenase and demonstrated the applicability of this proposed MIC biomarker assay in industrial settings.

## RESULTS

### Methanogenic archaeal communities on carbon steel pipeline walls

Maintenance of oil field pipelines involves the periodic use of devices with blades or brushes (termed ‘pig’, pipeline intervention gadget) to mechanically clean out rust, wax, scale and biofilm (collectively termed ‘pig debris’). This practice offers the opportunity to access steel-associated microorganisms from pressurized hydrocarbon pipelines. We obtained pig debris from nine different pipelines in seven geographically distinct oil field operations in North America, East Asia and West Africa to survey the prevalence of methanogenic archaea (Fig. 1). Estimated 16S rRNA gene abundances of 1.6 · 10^7^ - 1.3 · 10^10^ gene copies per g of pig debris suggested moderately dense microbial colonization of pipe wall-associated solids. Archaeal sequences accounted for 0.2 – 21.6% of total reads, and most of the detected archaea were classified as members of methanogenic archaeal families (0.2 – 9.3% of total reads). Members of the *Methanococcaceae*, *Methanobacteriaceae*, and *Methanosarcinaceae* have been implicated in MIC through laboratory investigations (13, 17, 19, 24). However, there was no apparent correlation between the types or abundances of methanogens, and the perceived severity of MIC in the surveyed pipelines (Fig. 1). For example, pipelines [A] – [C] had no indications of active internal corrosion (MIC or otherwise) in recent in-line inspections (ILI, data not shown) despite the presence of microorganisms affiliating with the abovementioned putatively corrosive archaeal families.

**Figure 1.**
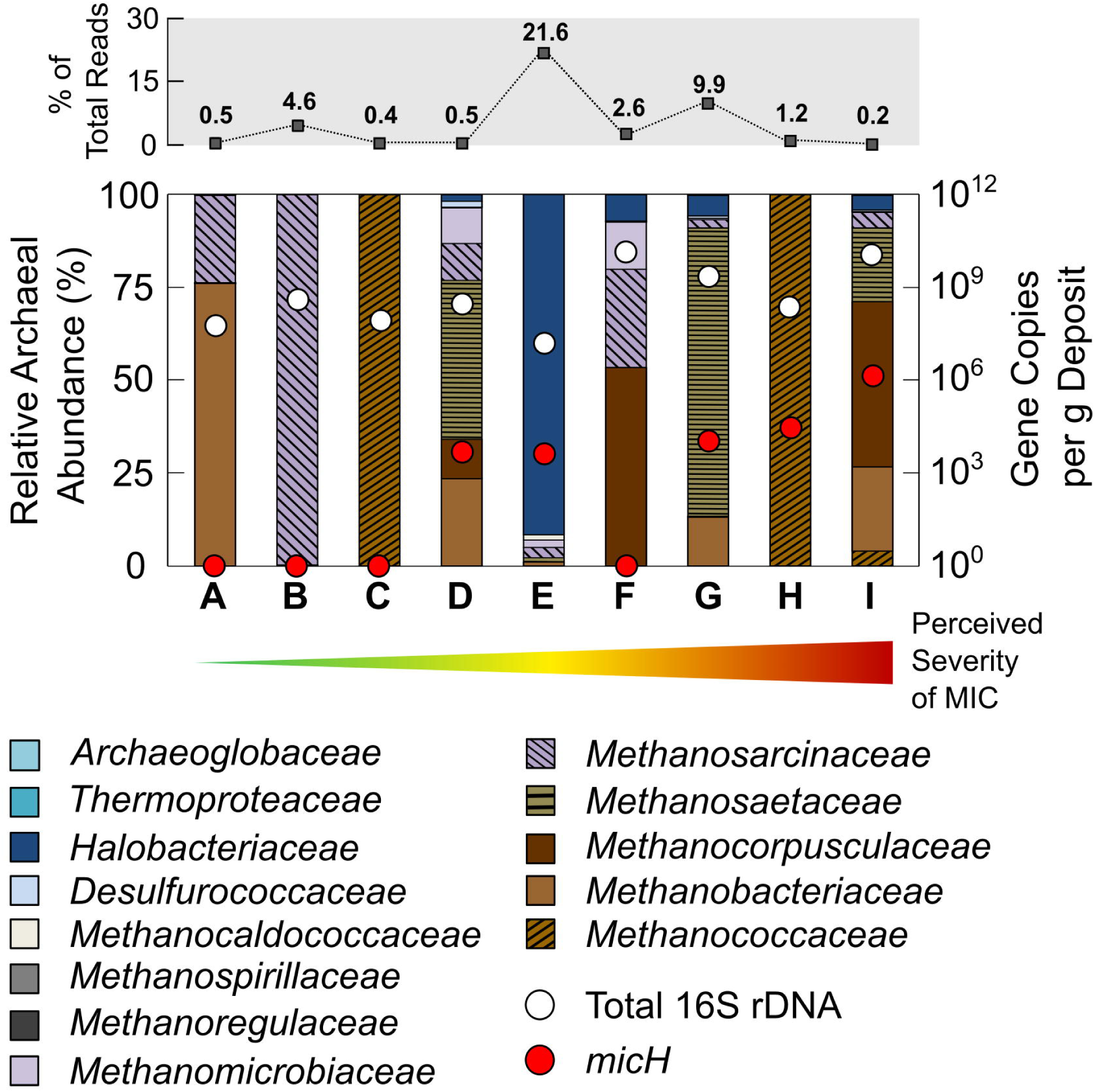
Archaeal community composition in solid samples (pig debris) collected from carbon steel pipelines. Archaeal sequences accounted for between 0.2% and 21.6% of all reads from 16S rRNA gene sequencing of DNA extracted from pipeline solids (top row). Total 16S rRNA gene numbers (archaeal + bacterial) and gene copy numbers of the proposed methanogenic MIC biomarker *micH* are also plotted for each sample. Pipelines (A – H) are ordered according to their perceived severity of MIC (increasing from left to right, as indicated by arrow). The perceived (qualitative) severity of MIC is based on inspection data (in-line inspection, ILI), field and laboratory studies, as well as operator experience. A: Offshore multi-phase pipeline (East Asia). B: Offshore crude transmission pipeline (U.S. Gulf of Mexico). C: Offshore multi-phase pipeline (U.S. West Coast). D: Onshore crude transmission pipeline (U.S. Gulf Cast). E: Onshore crude transmission pipeline (U.S. West Coast). F: Onshore produced water pipeline (U.S. Midwest). G: Offshore multi-phase pipeline (Nigeria). H: Offshore multi-phase pipeline (U.S. West Coast). I: Offshore multi-phase pipeline (Nigeria).

### Assessment of MIC under mesophilic and thermophilic conditions

The detection of methanogenic archaea in all surveyed pipelines necessitated a more nuanced understanding of their potential role in corrosion; a question that was approached through laboratory cultivation. We hence obtained anoxic production fluids from a West African oil field with a history of MIC (Fig. 1, pipelines [G] and [I]). Carbon steel infrastructure typically spans broad temperature gradients within the same oil field due to hot production fluids from reservoirs (often >> 60°C) cooling as they pass through networks of pipelines, pressure vessels and piping. Fluids from subsea pipeline [I] had an average temperature of 32°C at the time of sampling. In the laboratory, the sulfate-free (< 20 μM) produced water was transferred into glass bottles with specially designed carbon steel coupon holders with rubber stoppers and an anoxic headspace of N_2_-CO_2_. Bottles were incubated at 32°C and 60°C on rotary shakers (75 rpm) for 13 weeks; under conditions that are most representative of process dead legs or idled pipelines (i.e. stagnant to very low flow). Indeed, methanogenic biofilms developed on carbon steel surfaces exposed to the sulfate-free produced water, as evidenced by accumulation of CH4 in the test bottles (Fig. 2A) and detection of high numbers of steel-attached methanogenic archaea through qPCR and 16S rRNA gene sequencing (Fig. 2A and B). Methanogens accounted for 17.0 – 54.4% of the steel-attached biofilms. Archaeal communities grown at 32°C showed high abundances of hydrogenotrophic (i.e. lithotrophic) *Methanobacterium* spp. and *Methanocalculus* spp., with minor fractions of *Methanococcus* spp. Incubation at higher temperatures (60°C) led to formation of biofilms with prominent fractions of hydrogenotrophic *Methanothermobacter* spp. and, at lower prevalence, the acetoclastic *Methanosaeta* spp. (Fig. 2B). Mesophilic bacterial communities (32°C) contained high fractions of *Syntrophomonas* spp., *Kosmotoga* spp. and *Halomonas* spp. that might have grown through fermentation of residual organic compounds in the produced water, thereby producing acetate, CO_2_ and H_2_ (26–28). The majority of the biofilm community in bottles incubated at 60°C consisted of mesophilic bacteria unlikely to grow at that temperature (Fig. 2C). However, some thermophilic bacteria such as *Thermotoga*, *Fervibacterium* and *Caldithrix* were detected and could have grown via a fermentative metabolism (29–31). Consistent with the lack of sulfate in these natural oil field waters, we only detected minor fractions of SRB in steel-attached biofilms (cumulatively < 0.6% per biofilm). The different incubation temperatures resulted in markedly different metal damage (Fig. 2A). Microbial activity had negligible impact on carbon steel at 60°C, with an increase in corrosion rates of approximately 40% compared to sterile controls, but a low average corrosion rate of 0.02 ± 0.01 mm Fe^0^ yr^−1^. Mesophilic microorganisms, on the other hand, accelerated corrosion by a factor of 12 and led to localized damage with features as deep as 399 μm developing over 3 months (Fig. 2A and Fig. S1). The National Association of Corrosion Engineers (NACE) categorizes corrosion rates of carbon steel coupons > 0.13 mm Fe^0^ yr^−1^ as *high* and > 0.25 mm Fe^0^ yr^−1^ as *severe* (NACE SP0775-2013). Interestingly, despite the profoundly different impacts on corrosion, biofilms grown at 32°C and 60°C contained similar numbers of active methanogens (Fig. 2A). This observation implied that intrinsic properties of the mesophilic biofilm community rather than the overall cell number or methanogenic activity were linked to the observed acceleration of corrosion kinetics.

**Figure 2.**
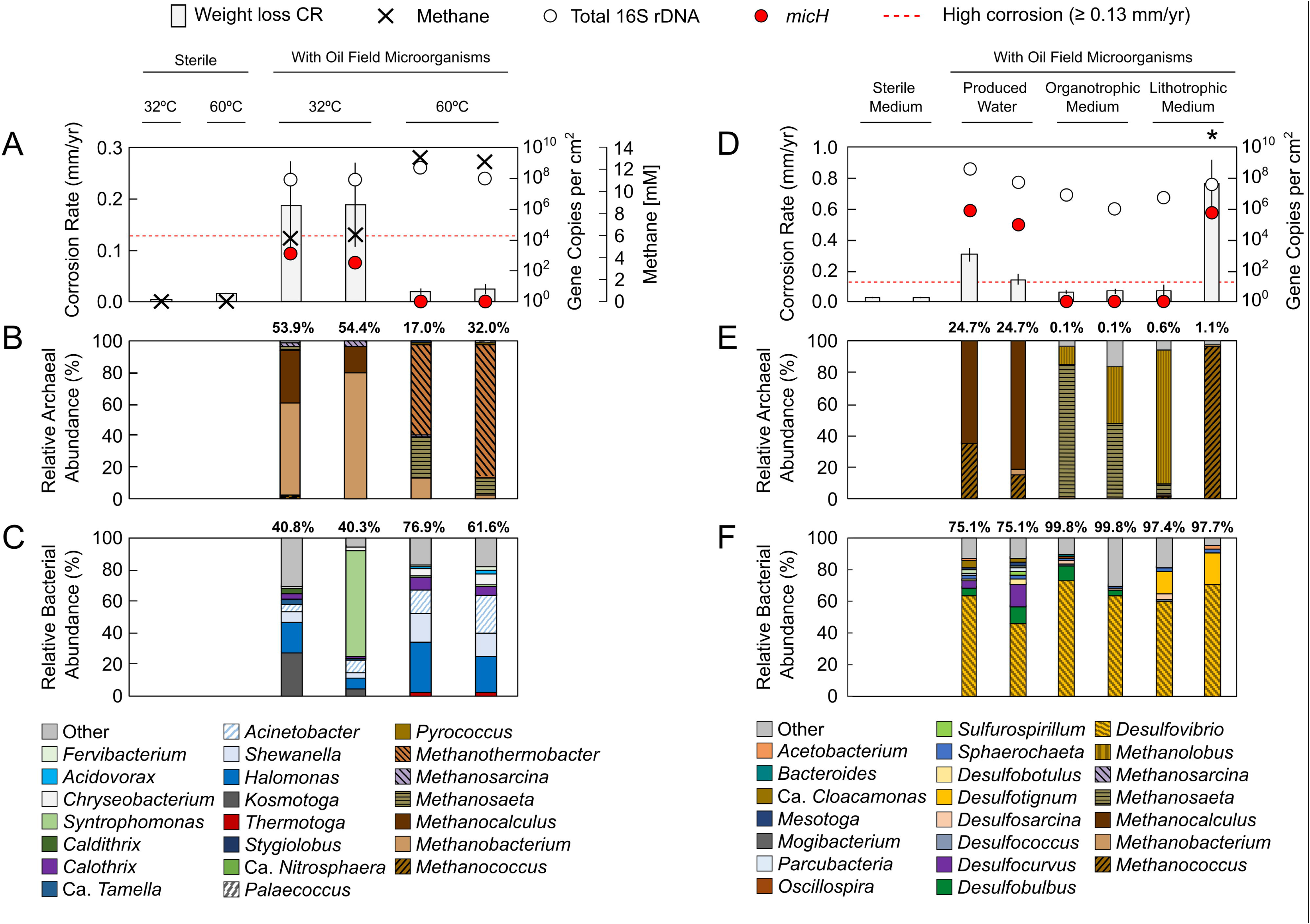
Growth of steel-attached oil field biofilms in bottle tests. A – C Tests with sulfate-free anoxic produced water as obtained from the West African oil field and incubated under mesophilic (32°C) and thermophilic (60°C) conditions. D – F Tests with sulfate-amended produced water, as well as synthetic produced water media (+ sulfate) in the presence or absence of propionate and acetate (organotrophic and lithotrophic conditions, respectively).). A and D Averaged weight loss corrosion rates (CR), total 16S rRNA gene (archaeal + bacterial) quantification and gene copy numbers of the proposed archaeal MIC biomarker *micH*. A threshold denoting technically relevant, high corrosion rates (≥ 0.13 mm Fe^0^ yr^−1^) is indicated by the dashed red line. Methane formation is depicted as final concentration in headspace after 3 months (A). Archaeal (B and E) and bacterial (C and F) community composition was assessed by 16S rRNA gene sequencing of DNA extracted from biofilms grown on carbon steel coupons. Archaeal sequences account for 0.1 – 54.4% and bacterial sequences for 40.3 – 99.8% of the total sequencing reads. The enrichment culture that was used for a subsequent lithotrophic corrosion kettle test is indicted by an asterisk.

### Identification of potential methanogenic corrosion mechanisms

Tsurumaru and colleagues recently linked the accelerated oxidation of Fe^0^ (i.e. corrosion) in cultures of *M. maripaludis* strain OS7 to a novel [NiFe] hydrogenase (24). A protein sequence comparison of the large subunit of this hydrogenase against the National Center for Biotechnology Information’s (NCBI) protein sequence database revealed that three sequenced members of the *Methanobacteriales* contain similar proteins (90.7 to 93.5% protein sequence identity), while all other deposited sequences showed ≤ 40% protein sequence identity (Table S1). The high degree of protein sequence resemblance suggested functional similarity within this protein cluster and prompted us to develop a specific qPCR assay. We developed a degenerated primer pair and Taqman™ probe (Table 1) to detect and quantify the gene of the large subunit of the [NiFe] hydrogenase (referred to here as *micH*) in strain OS7 and the three other methanogenic strains. This assay allows for probing of mixed microbial biofilms for their ability to carry out a proven microbial corrosion mechanism, i.e. the enzymatic catalysis of Fe^0^ oxidation via an extracellular hydrogenase. Intriguingly, we could detect 3.3·10^2^ and 1.6·10^3^ *micH* gene copies per cm^2^ in corrosive biofilms at 32°C while the *micH* gene was undetectable in non-corrosive biofilms grown at 60°C (Fig. 2A). This raised the possibility that mesophilic, lithotrophic methanogens, likely of the genera *Methanococcus* or *Methanobacterium*, had caused the observed high rates of corrosion.

**Table 1.**
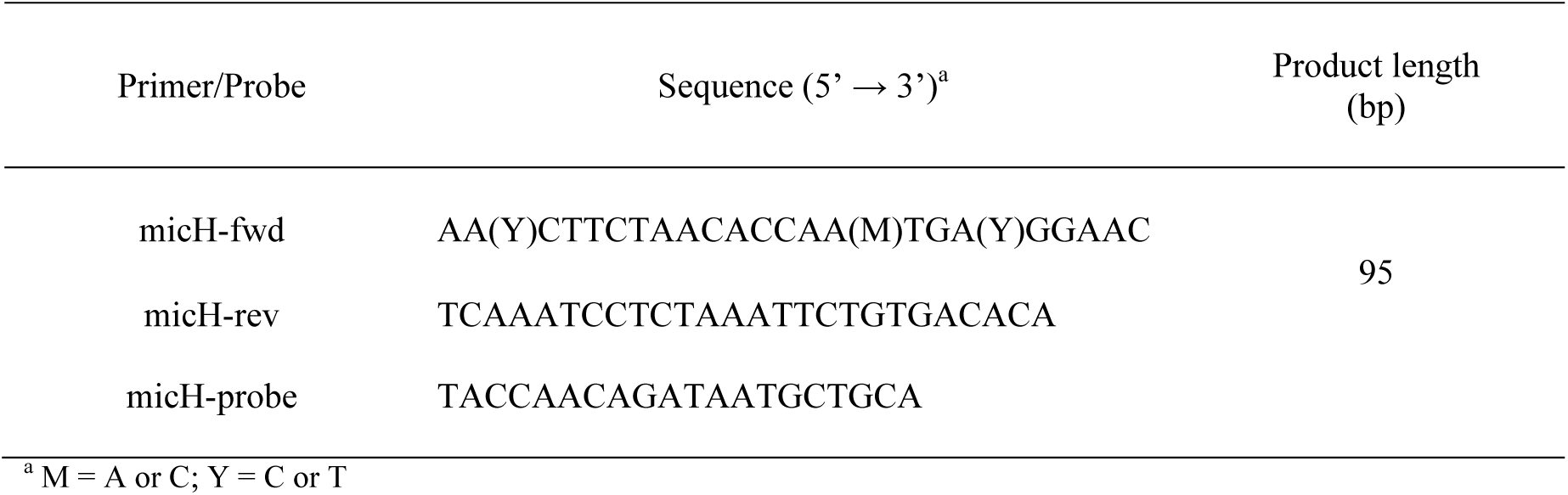
Sequences of degenerate primer pair and probe used in qPCR assays to detect the proposed methanogenic MIC biomarker *micH*.

### Enrichment of organotrophic and lithotrophic oil field communities

We sought to reproduce our observations and collected a new batch of production fluids from the same West African oil field. Traces of H_2_S (approx. 100 ppm) measured in the gas phase of the sample collection drums upon arrival in our laboratories indicated sulfate reduction, so we amended the next tranche of corrosion bottle tests with sulfate to allow for the growth of SRB. Again, highly corrosive mesophilic biofilms developed in these produced waters and putatively corrosive *Methanococcus* spp., as well as *Methanocalculus* spp. were detected in large fractions (Figs. 2D and 2E). Furthermore, these biofilms contained *micH* at even higher numbers than previous cultures with 1.1 · 10^5^ – 7.4 · 10^5^ gene copies per cm^2^ (Fig. 2D).

The use of actual produced waters for laboratory studies offers the opportunity to closely simulate oil field MIC in laboratory settings. However, it also complicates data interpretation, since the presence of complex organic matter can allow for a multitude of fermentative and oxidative metabolisms. We therefore dissected experimental conditions by working with defined cultivation media which were modelled after the major ion composition of the original produced water (+ sulfate). Organotrophic and lithotrophic enrichment cultures containing steel coupons were set up with or without addition of organic acids (12.1 mM acetate and 1.6 mM propionate), respectively. This reduced the number of potential biological transformations and, in the case of tests without the organic acids, limited growth to those microorganisms capable of utilizing metallic iron (Fe^0^) as an electron donor. Transfer of produced water as inoculum (0.5% vol/vol) into synthetic media reduced microbial corrosion rates to moderate rates (< 0.08 mm Fe^0^ yr^−1^) in all but one of the bottle tests (Fig. 2D). Remarkably, this lithotrophic culture caused corrosion at severe rates of 0.76 ± 0.15 mm Fe^0^ yr^−1^, with up to 370 μm deep corrosion features developing on carbon steel coupons over 3 months (Fig. 2D and Fig. S2). All biofilms in the synthetic produced water media were numerically dominated by sulfate-reducing bacteria, in particular *Desulfovibrio* spp., *Desulfobulbus* spp. and *Desulfotignum* spp. (Fig. 2F). The distinguishing feature of the severely corrosive lithotrophic biofilm, however, appeared to be the enrichment of a sub-population of *Methanococcus* spp., and the detection of the special [NiFe] hydrogenase *micH* (Figs. 2D and 2E).

### Corrosion under simulated pipeline condition

The preceding bottle tests demonstrated the potential for severe MIC in stagnant water bodies within infrastructure of the investigated oil field. However, the pipelines that transport multi-phasic fluids (e.g. from offshore production wells) usually experience high flow and, as a result, wall shear stresses, which greatly increase the rates of acid gas corrosion resulting from CO_2_ (32). Likewise, biofilm formation can be affected by the physical stresses imposed by flow (33).

We therefore evaluated corrosion in customized test reactors (corrosion kettles) that allow for close simulation of pipeline conditions by controlling shear stress, pH and acid gas composition (Fig. 3A). First, a test with sterile synthetic produced water was performed to establish the baseline corrosion from CO_2_. These tests revealed that abiotic conditions alone could cause severe corrosion of 0.34 ± 0.01 mm Fe^0^ yr^−1^ (Fig. 3B and Fig. S3) under conditions that model pipelines [G] and [I].

**Figure 3.**
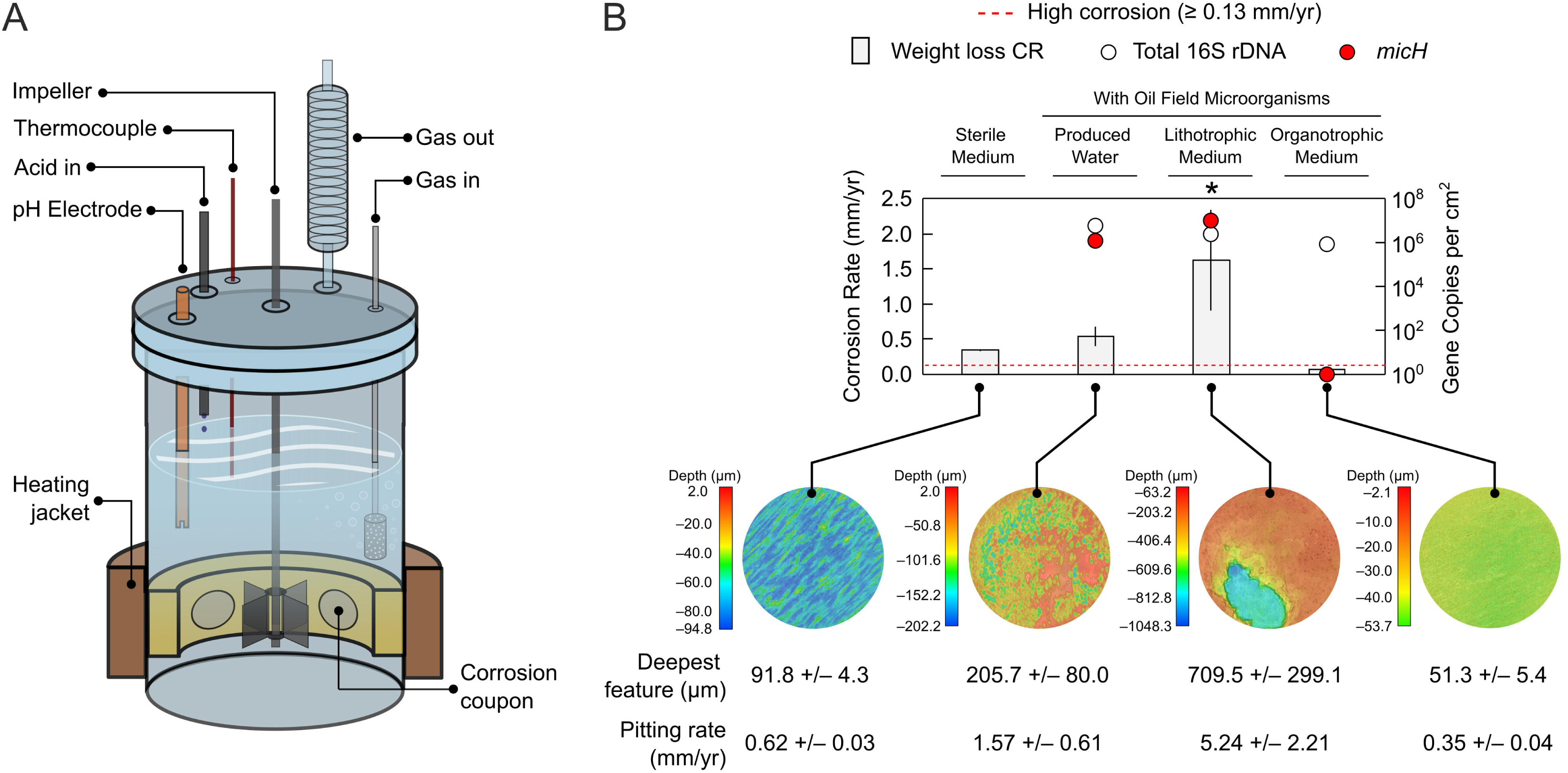
Growth of steel-attached oil field biofilms in corrosion kettle tests. A) Schematic of customized corrosion kettles for the study of CO_2_ corrosion and MIC under pipeline-simulating conditions (flow and acid-gas purging). B) Averaged weight loss corrosion rates (CR) in anoxic kettle tests containing oil field microorganisms in West African produced water, as well as synthetic produced water media with or without addition of the organic electron donors acetate and propionate (organotrophic and lithotrophic conditions, respectively). A sterile control test quantifying CO_2_ corrosion was also included. Total 16S rRNA gene (archaeal + bacterial) quantification and gene copy numbers of the proposed archaeal MIC biomarker *micH* in coupon biofilms are plotted for each test. A threshold denoting technically relevant, high corrosion rates (≥ 0.13 mm Fe^0^ yr^−1^) is indicated by the dashed red line. False color images show the topography of coupons at the end of the experiments after removal of corrosion products. The average maximal feature depth from four steel coupons per test and corresponding linearly extrapolated pitting corrosion rates are provided in the sub-panel. The asterisk indicates the test selected for metagenomic analysis.

Still, inclusion of oil field microorganisms in these tests demonstrated the pronounced biotic component of pipeline corrosion under the studied conditions; a second tests with actual produced water increased averaged corrosion rates beyond the CO_2_ baseline to 0.54 ± 0.14 mm Fe^0^ yr^−1^ (Fig. 3B), and resulted in irregular and localized pitting corrosion at extrapolated rates of 1.57 ± 0.61 mm yr^−1^ (Fig. 3B and Fig. S3). Operation of a third corrosion kettle with lithotrophic produced water medium inoculated with a pre-enriched Fe^0^-utilizing culture (asterisk in Fig. 2D) resulted in even more aggressive metal damage. This culture catalyzed iron oxidation at rates of 1.62 ± 0.72 mm Fe^0^ yr^−1^, leading to macroscopic localized corrosion features with an extrapolated growth rate of 5.24 ± 2.21 mm yr^−1^ over 7 weeks (Fig. 3C). Microbially catalyzed metal oxidation at these rates, if left unmitigated, would profoundly challenge pipeline integrity.

High copy numbers of *micH* (> 10^6^ copies per cm^2^) and high relative fractions of 28% and 42% of *Methanococcus* spp. were detected in biofilms grown in corrosion kettles operated with actual (organotrophic) and synthetic (lithotrophic) produced water, respectively (Fig. 3B and Fig. S4). Other dominant microorganisms detected in corrosive biofilms in both reactors included *Desulfovibrio* spp. (Fig. S4).

A fourth corrosion kettle was set up with synthetic produced water medium containing organic acids and was inoculated with an organotrophic subculture from the West African oil field. This test did not produce the previously observed high corrosion rates, despite formation of similar biofilms as measured by 16S rRNA gene abundance (Fig. 3B). Corrosion of coupons underneath these biofilms containing *Desulfovibrio, Desulfobulbus* and *Methanolobus* species (Fig. S4) was considerably lower than the sterile control, at 0.073 ± 0.001 mm Fe^0^ yr^−1^ and resulted in rather uniform metal loss (Fig. 3B and Fig. S3). This may be an indication that the biofilm limited CO_2_ corrosion by forming a passivating barrier, a phenomenon seen before in laboratory tests (Kip and van Veen 2015 and references within). Notably, this ‘protective’ biofilm did not contain any *Methanococcus* spp. or *Methanobacterium* spp. and tested negative for *micH* (Figs. 3B and S5).

### Genomes recovered from a severely corroded carbon steel coupon

Shotgun metagenome analysis can be a useful tool to probe for blueprints of biochemical reactions in mixed microbial communities. We wanted to further investigate the molecular mechanisms of MIC seen in this study, and hence extracted and sequenced DNA of a biofilm that was i) associated with severe corrosion, ii) contained high fractions of potential culprit organisms such as *Methanococcus* spp. and *Desulfovibrio* spp. and iii) was grown under pipeline-like conditions (see asterisk in Fig. 3B for selected sample).

We were able to obtain six near complete metagenome-assembled genomes (MAGs) from this sample (Table 2). Taxonomic classification by protein marker analysis indicated that one of the MAGs affiliated closest with *Methanococcus maripaludis*. The *micH* gene in *M. maripaludis* strain OS7 is co-located on a 12 kb ‘MIC island’ with genes hypothesized to be essential for the functioning and secretion of this hydrogenase (24). A sequence similarity search revealed that a *micH* homolog was indeed present in one of the assembled contigs of the *M. maripaludis* MAG. Furthermore, we discovered that the genetic arrangement of the ‘MIC island’ in this MAG and strain OS7 was in fact identical (Fig. 4). Notably, the genetic island contained genes for both subunits of the putative [NiFe] hydrogenase, as well as the twin-arginine translocation system for secretion of the mature hydrogenase in strain OS7 (24). Protein sequence comparison further confirmed a high resemblance (98.8 – 100% amino acid sequence identity) between both clusters (Fig. 4).

**Table 2.**
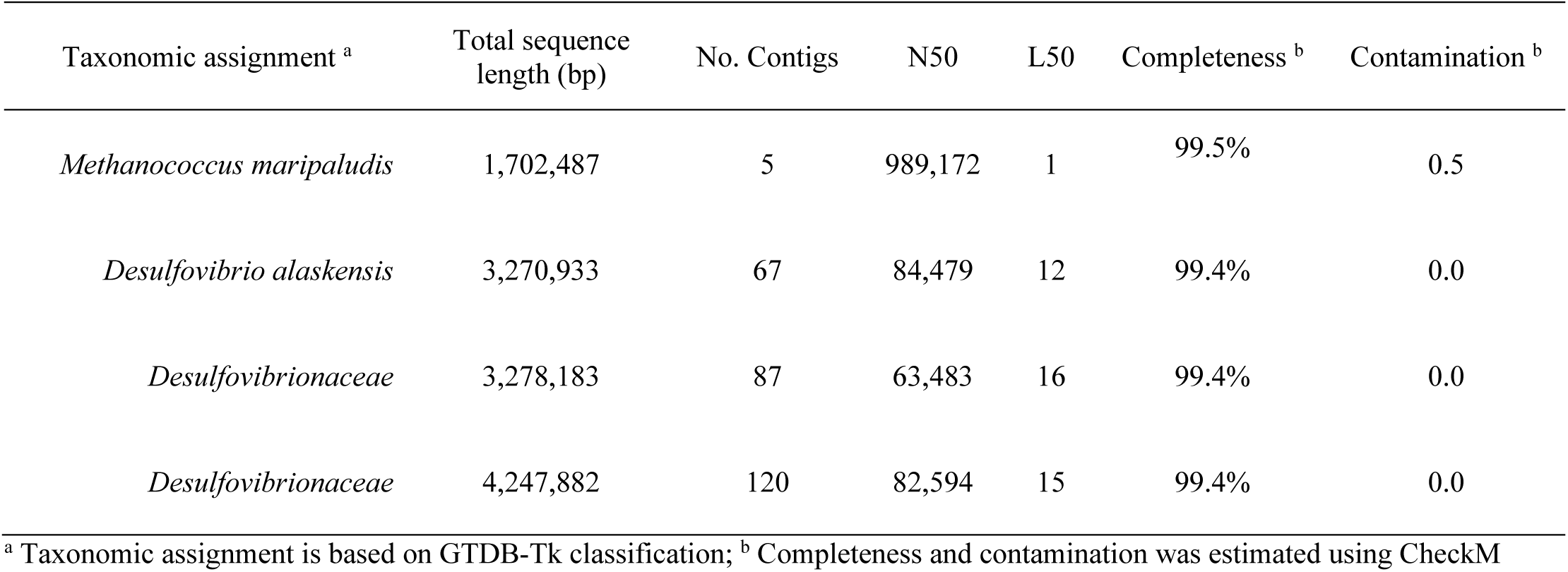
Metagenome-assembled genomes (MAG) of methanogenic and sulfate-reducing microorganisms in severely corrosive biofilm.

| Taxonomic assignment <sup>a</sup> | Total sequence length (bp) | No. Contigs | N50 | L50 | Completeness <sup>b</sup> | Contamination <sup>b</sup> |
| --- | --- | --- | --- | --- | --- | --- |
| *Methanococcus maripaludis* | 1,702,487 | 5 | 989,172 | 1 | 99.5% | 0.5 |
| *Desulfovibrio alaskensis* | 3,270,933 | 67 | 84,479 | 12 | 99.4% | 0.0 |
| *Desulfovibrionaceae* | 3,278,183 | 87 | 63,483 | 16 | 99.4% | 0.0 |
| *Desulfovibrionaceae* | 4,247,882 | 120 | 82,594 | 15 | 99.4% | 0.0 |

<sup>a</sup> Taxonomic assignment is based on GTDB-Tk classification; <sup>b</sup> Completeness and contamination was estimated using CheckM

**Figure 4.**
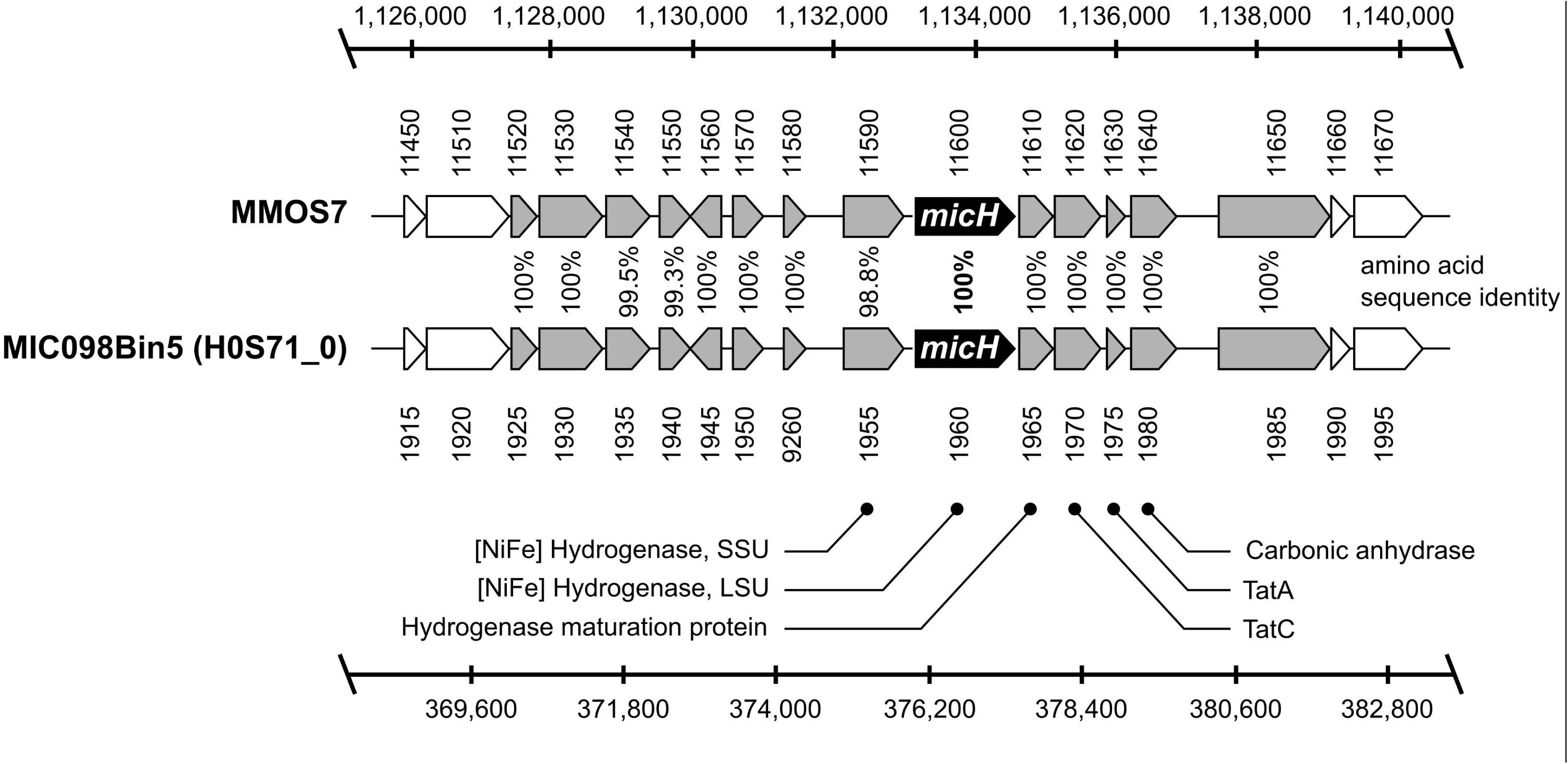
Comparison of the genetic arrangement of the ‘MIC island’ in *Methanococcus maripaludis* strain OS7 (MMOS7) and the metagenome-assembled genome (MIC098Bin5) of a *M. maripaludis* strain grown on severely corroded steel surfaces. The scale bars indicate the nucleotide position in the individual genomes.

Traditionally, SRB have been viewed as prime suspects of MIC (7). While all SRB can produce corrosive H_2_S under suitable organotrophic growth conditions, some specialized lithotrophic SRB strains have recently received much attention as they can cause particularly severe corrosion through a more direct mechanism (8, 9, 13). It had been proposed that these organisms oxidize metal through direct electron uptake via outer-membrane c-type cytochromes (OMCs, (13)), with recent evidence supporting this hypothesis in *D. ferrophilus* strain IS5 (11). In this study, we obtained three near complete *Desulfovibrionaceae* genomes (Table 2). A reliable taxonomic classification was possible for one genome, which was assigned to a strain of *Desulfovibrio alaskensis*. However, despite presence of three near complete *Desulfovibrionaceae* genomes, no evidence for a homologous OMC system to the one detected in strain IS5 was found in this metagenomic dataset. This suggest that a direct electron uptake via extracellular c-type cytochromes may not have contributed to MIC in our experiments.

The detection of *micH* in a metagenome-assembled genome from a highly corrosive biofilm further strengthened the applicability of this gene as a potential marker for MIC. We confirmed the presence of *micH* (up to 1.6·10^6^ gene copies per g) in pig debris from the West African oil field (pipelines [G] and [I] in Fig. 1). Intriguingly, we also detected 3.9·10^3^ – 2.8·10^4^ *micH* gene copies per g in three geographically separated pipelines in North America (Fig. 1).

## DISCUSSION

Methanogenic archaea are commonplace members of the oil field microbiome (16, 17, 34, 35). However, unlike SRB which produce corrosive sulfides as a metabolic product, methanogenic archaea are not intrinsically corrosive (24, 36); they oxidize molecular hydrogen or a limited spectrum of small organic compounds and produce chemically inert CH4. We surveyed surface-associated deposits from nine pipelines in seven geographically distinct oil field operations and detected methanogenic archaea in all instances, despite vast differences in the perceived severity of MIC (Fig. 1). For example, methanogens affiliated with the *Methanobacteriaceae*, *Methanococcaceae* and *Methanosarcinaceae* were detected in steel-associated solids from pipelines [A] - [C]. Members of these archaeal families have been implicated in MIC through laboratory studies (13, 17, 19, 24), yet recent inspections of the pipelines [A] – [C] did not reveal any indications of internal corrosion. Similarly, biofilms containing *Methanosaeta* spp., *Methanobacterium* spp., *Methanolobus* spp. and *Methanothermobacterium* spp. caused only negligible corrosion (< 0.08 mm Fe^0^ yr^−1^) in laboratory reactors that simulated oil field conditions (Figs. 2, 3 and S5). These data highlight that the mere presence of methanogenic biofilms on steel does not entail corrosion.

Corrosion of metallic iron in the absence of oxygen can be depicted as follows:

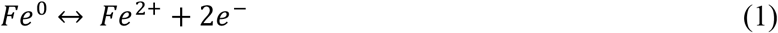

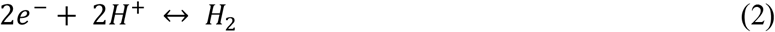

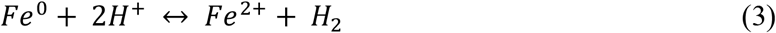

Where the anodic oxidation of iron (reaction 1) is coupled to the cathodic reduction of a suitable oxidant, in anoxic environments usually H^+^ (reaction 2), so that the net reaction involves formation of molecular H_2_ (reaction 3). Reaction 2 is kinetically impeded and as a result the corrosion of steel is slow in anoxic environment at near-neutral pH (9, 37, 38). However, corrosion in (slightly) acidic oil field waters can be pronounced in the presence of weak acids, as loosely bound protons from carbonic acid, organic acids or sulfide dramatically increase the availability of H^+^ ions as oxidants at the steel surface, particularly in high flow scenarios (39).

Historic models of MIC had postulated that the consumption of cathodic molecular hydrogen by hydrogenotrophic microorganisms (incl. methanogens) would accelerate corrosion (40–42). However, this view has been disputed based on kinetic and thermodynamic considerations (8, 9, 43), as well as through copious experimental observations (8, 13, 14, 24, 36). However, while the majority of tested hydrogenotrophic methanogenic archaea does not influence steel oxidation to any significant extent, some strains have shown a pronounced impact on corrosion kinetics in laboratory experiments (13, 14, 24, 36). Recently evidence has emerged that some pure cultures of methanogenic archaea accelerate corrosion when they produce extracellular hydrogenases that catalyze reaction 2 on steel surfaces (23, 24).

### Mesophilic methanogens in highly corrosive oil field biofilms contain a special MIC hydrogenase

In this study we reproduced oil field corrosion using fluids from an offshore production site in West Africa. Formation of mesophilic biofilms considerably increased corrosion rates over sterile controls, both in *de facto* stagnant bottle test, as well as in corrosion kettles simulating flow and field-like partial pressures of CO_2_ (Figs. 2 and 3). Technically highly relevant MIC rates (0.15 – 1.62 mm Fe^0^ yr^−1^) and severe pitting were observed in several of these bioreactors. By using specially formulated synthetic produced water media we could further demonstrate that the observed Fe^0^ oxidation was in fact a lithotrophic process. Some hydrogenotrophic *Methanococcus* spp. and *Methanobacterium* spp. have been shown to effectively utilize Fe^0^ and were found in substantial numbers (> 3.7 · 10^5^ 16S rRNA gene copies per cm^2^) in highly corrosive biofilms (Figs. 2 and 3; Fig. S5B). However, both genera were also present in nearly all other (non-corrosive) biofilms, confirming that hydrogenotrophy itself does not accelerate corrosion kinetics.

Tsurumaru and colleagues recently discovered that only a few known strains of *M. maripaludis* and *Methanobacterium* spp. encode for a unique hydrogenase that catalyzes the reduction of H^+^ to H_2_ on Fe^0^ granules and thereby considerably accelerates iron oxidation (24). The apparent high conservation amongst proteins from members of *Methanococcaceae* and *Methanobacteriaceae* allowed us to develop a specific qPCR assay targeting the gene of the large subunit, *micH*, of this hydrogenase. This assay did in fact distinguish biofilms causing technically relevant corrosion (> 0.13 mm Fe^0^ yr^−1^) from those that did not (< 0.08 mm Fe^0^ yr^−1^) in our experiments with actual and synthetic produced waters (Fig. S5B, Figs. 2 and 3). *M. maripaludis* strain OS7 harbors this unique hydrogenase (*micH*) on a 12 kbp long genetic island along with several other genes assumed to be essential for a fully functional extracellular enzyme (24). We were able to retrieve a near complete genome of a *Methanococcus maripaludis* strain that carried *micH* in an identical genetic arrangement as strain OS7 (Fig. 4). This indicated that *micH* was associated with a corrosive *Methanococcus maripaludis* strain in our laboratory tests. Indeed, corrosive biofilms consistently contained both, methanococcal 16S rDNA and the *micH* gene (Fig. S5B).

There was no linear correlation between the extent of corrosion and copy numbers of *micH* (Fig. S5). High corrosion rates were observed on coupons containing as few as 3.3·10^2^ gene copies per cm^2^ at the time of biofilm collection (Fig. 2A). It should be noted though that corrosion rates were averaged from metal loss over the entire experimental period (e.g. 3 months), while quantification and characterization of target genes in biofilms was only performed at test takedown and hence only represents endpoint conditions. Methanogens precipitate siderite (FeCO_3_) which can passivate steel surfaces (18, 44) and could have buried initially corrosive methanogens in some tests, possibly allowing for the degradation of ‘forensic evidence’. This could explain low copy numbers in some highly corrosive biofilms. More importantly though, we do not envision gene copy numbers of *micH* to correlate with the final abundance of the actual enzyme that catalyzes iron oxidation. Still, *micH* in biofilms served as a binary genetic marker for technically relevant microbial corrosion in our tests with West African produced waters.

Methanogenic archaea constituted only a part, and in some instances a minor part of the biofilm communities that developed under simulated oil field conditions. In addition to organic compounds in the produced waters and the organotrophic synthetic produced water medium (e.g. acetate and propionate), also small quantities of cathodic hydrogen from abiotic corrosion according to reaction 3 will have served as a readily available electron donor, e.g. for sulfate-reducing bacteria. It is tempting to speculate that enzymatically catalyzed H_2_ formation from extracellular hydrogenases could have fed additional electron donors into the biofilm community. A potential role of electroactive H_2_-forming microorganisms as ‘ecosystem engineers’ has been proposed elsewhere (45).

### The role of sulfate-reducing bacteria in corrosion

The importance of sulfate-reducing bacteria in microbial corrosion has long been established ((7) and references therein) and the underlying mechanisms as well as their regulation are still an active area of research (11, 46–48). While we cannot exclude contribution of SRB to corrosion, we inferred that they had likely played a minor role in our experiments. Firstly, *Desulfovibrio sp*. were found in high abundances in all biofilms under sulfate-reducing conditions, but high or severe corrosion only occurred in the presence of *micH*-positive methanogens (Figs. 2D and 3B; Fig. S5). Secondly, high microbial corrosion rates were also observed in sulfate-free produced water (Fig. 2A). Thirdly, only few lithotrophic SRB isolates have been demonstrated to cause corrosion to an extent similar to what was observed in this study (7, 8); the extracellular multiheme c-type cytochromes that have been linked to direct electron uptake in those corrosive strains (11) were absent in the metagenomic dataset. The low to moderate corrosion rates in the absence of *micH*-positive *Methanococcus* spp. agree with corrosion rates documented for sulfidogenic cultures of *D. alaskensis* (≤ 0.1 mm Fe^0^ yr^−1^; (22, 48, 49)). We cannot, however, exclude that hitherto unknown enzymatic mechanisms may have affected corrosion in our experiments.

### Proposed use of *micH* as a genetic marker for methanogenic MIC

Microbial corrosion of oil field infrastructure is difficult to detect and monitor, in large part because the responsible microorganisms and underlying mechanisms remain enigmatic (50). For example, the mere presence of large numbers of methanogenic archaea in pipeline-associated debris or laboratory-grown biofilms proved to be a poor indicator of MIC in this study (Fig. 1 and Fig. S5). A deeper mechanistic understanding of MIC in methanogens and other environmental microorganisms offers a route to a more granular analysis and interpretation of microbiological monitoring data in oil fields and other industrial settings. Enzymatic acceleration of corrosion via hydrogenases is one such potentially relevant corrosion mechanism. This was recently evidenced through careful experimentation with pure cultures of methanogenic archaea (24) and demonstrated in this study with mixed microbial communities in bioreactors simulating oil field corrosion. We designed an assay to quantify the gene encoding for an extracellular hydrogenase (*micH*) in methanogens and demonstrated its ability to serve as a binary marker for MIC in experiments with oil field cultures grown under stagnant and pipeline flow-simulating conditions (Fig. S5B). The proposed MIC biomarker was detected in high concentrations (up to 1.6·10^6^ gene copies per g) in pipeline-associated solids from a West African oil field with a history of MIC (pipeline [I] in Fig. 1). Interestingly, we also detected *micH* in similar solids from pipelines on the West Coast and Gulf Coast of the United States (Fig. 1). The genetic island carrying this gene was first detected in *M. maripaludis* strain OS7, which originated from an underground oil storage tank in Japan and was further detected in the genome of *Methanobacterium congolense* Buetzberg by the same group (24). Strain Buetzberg was originally isolated from a biogas plant in Germany (51). The fact that we were able to reconstruct a near identical ‘MIC island’ from a West African oil field strain of *M. maripaludis* (Fig. 4) suggests an extremely high level of conservation of the ‘MIC island’ and presence of these corrosive methanogens in industrial facilities across four continents. Future work may focus on developing a better understanding for the quantitative interpretation of *micH* in solid and liquid samples to enable early detection and mitigation of severe cases of microbially influenced corrosion in industrial settings.

## MATERIALS AND METHODS

### Collection of solid samples (pig debris) from pipelines

Solids from nine carbon steel pipelines were collected between July 2014 and June 2017 to study the microbiomes within hydrocarbon-transporting infrastructure. Subsea pipelines [G] and [I] transport fluids from one oil field offshore Nigeria. This field produces oil and sweet natural gas (i.e. no H_2_S). Subsea pipelines [C] and [H] transport fluids from an offshore oil field on the U.S. West Coast. The other five pipelines each carry fluids from separate oil and gas fields on the U.S. West Coast [E], Midwest [F] and Gulf Coast [B, D], as well as from East Asia (A). Samples were collected following the (internal) mechanical cleaning of these pipelines with maintenance pigs that remove and push out surface-associated and settled debris. These solid samples (collected in the pig trap) were carefully transferred into sterile containers, immediately refrigerated with ice packs and frozen in common household freezers within 4 hours of collection. Samples were then shipped frozen to Houston (USA) where they were stored at −80°C until further analysis.

### Bottle corrosion testing with original produced water

Production fluids (oil with associated water and gas) were collected at a pipeline outlet [I] in a large oil field offshore Nigeria in December 2014. The fluids were filled into an internally PTFE-lined steel drum (approx. 20 L) all the way to the brim and capped gas-tight. In the laboratory, the content of the gas-tight drum was purged with N_2_-CO_2_ (79:21) to remove any biogenic H_2_S that had formed in the drum during transit (8 weeks). Thereafter, anoxic and sterile technique was used to transfer the gravity-separated produced water into 1 L glass bottles (600 mL each; pH 6.6 – pH 6.7) with butyl rubber stoppers under a N_2_-CO_2_ (79:21) headspace. For sterile incubations, the capped produced water bottles were autoclaved (anaerobically) at 121°C for 20 minutes. Three sterilized carbon steel coupons (3 x 1.25 cm^2^) in a coupon holder made of polyether ether ketone (PEEK) were added to each incubation (see Lahme et al., 2019 for details). Bottles were placed on rotary shakers (75 rpm) and incubated at 32°C or 60°C for 13 weeks.

In March 2016, a second batch of production fluids was collected at a location further downstream from the same pipeline [I], shipped to the laboratory (transit time: 5 weeks) and processed as described above. Here, bottles with anoxic produced water (pH 6.6) were amended with 1 mM sodium sulfate (Na_2_SO_4_) and incubated at 32°C for 6 weeks.

### Bottle corrosion testing with synthetic produced water medium

To study microbial corrosion under more defined conditions, bottle tests with synthetic produced water medium were set up. The medium was modeled after the Nigerian produced water (collected in March 2016 referred to below as PW-2016) and contained (all Sigma-Aldrich, reagent-grade chemical in RO-DI water): NaCl (15.31 g/L), MgCl_2_·6H_2_O (0.562 g/L), CaCl_2_·2H_2_O (0.434 g/L), KCl (0.121 g/L) and NaBr (0.064 g/L). Additionally, Na_2_SO_4_ (1.48 g/L) was added to allow for microbial sulfidogenesis. The brine was aliquoted (600 mL) into 1 L glass bottles, purged with N_2_-CO_2_ (79:21), sealed with butyl rubber stoppers and autoclaved. After cooling, 24 mL/L of a sterile 1M NaHCO_3_ stock solution (equilibrated with pure CO_2_) was added. Subsequently, the brines were supplemented with thiamine, a vitamin mixture, riboflavin, vitamin B12, a trace element mixture, selenite and tungstate, potassium phosphate and ammonium chloride from sterile stock solutions (1 ml/L each), prepared as described previously (52). For organotrophic test conditions, the medium was further supplemented with sodium acetate (NaCH_3_CO_2_; 0.996 g/L) and sodium propionate (NaC_3_H_5_O_2_; 0.157 g/L). The pH of the synthetic produced water medium ranged from pH 6.6 to pH 6.7. Sterile carbon steel coupons in PEEK coupon holders were added as described above, bottles were inoculated with 0.5% (vol/vol) of the original field produced water and then incubated on rotary shakers (75 rpm) at 32°C for 11 weeks.

### Kettle corrosion testing

Customized corrosion kettles were used to simulate the influence of pipeline flow on CO_2_ corrosion and MIC (Fig. 3A). These reactors contained inward-facing corrosion coupons that were flush-mounted in an inert coupon holder cage (PEEK) and thus arranged around a central impeller with Rushton blades. The rotational speed was set to 500 rpm to correspond to an average coupon wall shear stress of 7 Pa (7 kg m^−1^ s^−2^), which was deemed representative of pipeline conditions based on multi-phase flow modeling previously performed at this field. The glass reactor containing 1.5 L of test brine was sealed air-tight, with continuous purging of N_2_-CO_2_ (79:21; equivalent to 3 psia CO_2_) through a diffusor (250 mL/min) and a gas outlet with refrigerated condenser tube to minimize loss of liquids via evaporation. Corrosion kettles were temperature-controlled at 32°C by means of a heating jacket with thermo-couple. Control of pH was achieved through titration with 1 M HCl via syringe pumps coupled to on-line pH monitoring and a set point of pH 6.6. All tests contained four carbon steel coupons (5 cm^2^ each) and were operated for 7 weeks, unless otherwise stated.

Prior to test setup, corrosion kettles were rigorously cleaned and then “de-contaminated” through liberal application of acetone. Sterile test conditions cannot be achieved in corrosion kettles. To minimize the growth of environmental contaminants, abiotic corrosion tests were amended with the antibiotics kanamycin (100 mg/L), chloramphenicol (25 mg/L) and tetracycline (20 mg/L), which did not affect acid gas corrosion rate or morphology in 7-day tests (Fig. S6). Furthermore, vitamins, trace elements, phosphate and ammonium were omitted from abiotic corrosion tests.

Corrosion testing was initially performed with the original PW-2016. This test included 1% (vol/vol) crude oil, was amended with sodium sulfate (10.4 mM; Na_2_SO_4_) and spiked with an Fe^0^-containing, sulfate-amended PW pre-enrichment as additional inoculum (3% vol/vol). Testing under lithoautotrophic conditions was performed in synthetic produced water medium (no acetate, propionate, or oil), inoculated with the first passage of a lithoautotrophic enrichment culture (0.5% vol/vol) obtained from PW-2016. Finally, kettle corrosion testing was also performed with synthetic produced water medium containing sodium acetate and sodium propionate as organic electron donors for microbial metabolism and growth. This test was inoculated with the third subculture of an organotrophic, Fe^0^-containing enrichment (0.5% vol/vol) obtained from PW-2016.

### Gas chromatography

Methane in gaseous samples (10 mL) was quantified using an Agilent 490 Micro gas chromatograph with thermal conductivity detector. Chromatography was performed on a heated Agilent PoraPlot U column (50°C, 22.1 psi, 10 m), using argon as the carrier gas. Injector and sample line temperature were set to 100°C.

### Weight loss corrosion analysis

API 5L X52 carbon steel coupons (≤ 0.26% C, ≤ 0.45% Si, ≤ 1.6% Mn, < 0.03% P, ≤ 0.03% S, balance Fe) were successively sonicated in hexane, acetone and methanol (5 min each), dried in a stream of air and placed under vacuum overnight. The weight of each coupon was measured three times on an analytical balance before placing into PEEK coupon holders. Coupon holders containing corrosion coupons were sterilized by soaking in ethanol (96%) for 10 minutes. Excess ethanol was dried with a stream of filtered N_2_ gas.

At test end, corrosion products and other deposits were dissolved from coupons in a warm (70°C) 1.8 N HCl solution containing 2% propargyl alcohol (1 min). Wet coupons were then neutralized in saturated calcium hydroxide solution (30 sec) and scrubbed with a nylon brush while being rinsed with deionized water. This procedure was performed twice. Coupons were then rinsed in acetone, dried under a stream of air and placed under vacuum overnight. To determine weight loss, each coupon was weighed three times. Corrosion rates were calculated by considering the weight loss, exposed surface area, steel density (7.87 g/cm^3^) and exposure time.

### Localized corrosion analysis

To visualize and measure localized corrosion, the topography of cleaned carbon steel coupons was mapped with a Keyence VR-3000K Wide-Area 3D profile measuring macroscope (Keyence Corp., Itasca, IL). Corrosion features (≥ 25 μm in depth) were quantified by referencing the original (pre-corrosion) surface height. Pitting corrosion rates, expressed as millimeter per year, were calculated by linear extrapolation.

### DNA sampling and extraction

Coupon holders were removed from bioreactors (bottles or corrosion kettles) and thoroughly rinsed with sterile phosphate buffered saline (PBS) solution to remove loose (planktonic) microorganisms. Steel-attached biofilms were then sampled using sterile swabs (minimum of two swabs per coupon) and frozen at −80°C.

Frozen samples (pig debris or biofilm swabs) were shipped on ice to Microbial Insights Inc. (Knoxville, TN) and DNA was extracted with the DNA Power Soil Total DNA Isolation kits (MO BIO Laboratories, Inc., Solana Beach, CA) according to manufacturer’s instructions. DNA quantity and purity were assessed with a NanoDrop^®^ ND-1000 spectrometer (Thermo Fisher Scientific, Waltham, MA).

### Quantitative polymerase chain reaction

Quantitative PCR was performed on a QuantStudio™ 12K Flex Real-Time PCR System using a Taqman™ Universal PCR Mastermix (Applied Biosystems, Grand Island, NY). For the quantification of eubacteria and archaea, primers and Taqman™ probes that target highly conserved regions of the 16S rRNA gene were used as described elsewhere (53, 54). Forward and reverse primers were obtained from Integrated DNA Technologies (Corralville, IA), while Taqman™ probes were synthesized by Thermo Fisher Scientific.

For the qPCR assay targeting the large subunit of the specific [NiFe] hydrogenase (here referred to as *micH*) found in highly corrosive methanogenic archaea ((24) and this study), a degenerated primer pair and Taqman™ probe were designed based on conserved regions in homologous genes (Table 1, Fig. S7). Appropriate primer pairs and probes were selected using the Primer Express™ software (Thermo Fisher Scientific) and initial specificity was assessed by BLASTN searches against NCBI’s nucleotide database (55). The qPCR reaction was performed in 20 μl using the Taqman™ Universal PCR Mastermix. Forward and reverse primers, as well as the probe were adjusted to a final concentration of 300 nM. For calibration and as positive control a synthetic fragment of the *micH* gene was used (Fig. S7 and S8). The qPCR reaction conditions were as follows: one cycle of 2 min at 50°C, followed by one cycle of 10 min at 95°C, 40 cycles of 15 s at 95°C and 1 min at 60°C. The specificity of the qPCR reaction was verified by PCR melt curve analysis and Sanger sequencing of final PCR products. In addition, potential cross reactions were excluded by running the newly developed qPCR assay on genome-sequenced cultures (Table S2) that lack the special [NiFe] hydrogenase (*micH*).

### 16S rDNA amplicon sequencing and analysis

16S rDNA amplicon sequencing was performed by GENEWIZ Inc. (South Plainfield, NJ). DNA quality was assessed by agarose gel electrophoresis and quantified on a Qubit 2.0 Fluorometer (Invitrogen, Carlsbad, CA). Barcoded amplicons covering the V3 to V5 region of the 16S rRNA gene were generated using GENEWIZ’s proprietary 16S MetaVx™ amplification protocols and library preparation kit. Sequencing libraries were evaluated on an Agilent 2100 Bioanalyzer (Agilent Technologies, Palo Alto, CA) and quantified on a Qubit Fluorometer as well as through qPCR. Paired-end sequencing (2× 250 bp) of multiplexed DNA libraries was performed on an Illumina MiSeq platform (Illumina, San Diego, CA).

The 16S rRNA data was analyzed using the QIIME pipeline (56). Joined reads were quality filtered, discarding reads with a mean quality score ≤ 20, a read length < 200 bp and reads containing any ambiguous bases. Chimeric sequences were detected using the UCHIME algorithm (https://drive5.com/uchime/uchime_download.html) and the Ribosomal Database Program (RDP) Gold Database (57). After removal of chimeric sequences, the remainder of sequences was assigned into operational taxonomic units (OTU) with the VSEARCH (v1.9.6) clustering program and the SILVA 119 database (58), pre-clustered at 97% identity. Taxonomic categories were assigned to all OTUs at a confidence threshold of 0.8 by the RDP classifier.

### Shotgun metagenome sequencing and analysis

Shotgun sequencing of extracted DNA was performed by GENEWIZ Inc. The quality of extracted DNA was assessed by agarose gel electrophoresis and quantified on a Qubit 2.0 Fluorometer. DNA libraries were prepared using the NEBNext^®^ Ultra™ DNA Library Prep Kit (New England BioLabs Inc., Ipswich, MA) according to manufacturer’s instructions. Quality of DNA libraries was evaluated using Agilent TapeStation and quantified using a Qubit Fluorometer. Libraries were quantified by qPCR and then loaded on to an Illumina HiSeq 4000 platform according to manufacturer’s recommendations and sequenced using the paired-end (2× 150 bp) configuration.

Subsequent bioinformatics analysis of paired-end sequencing reads was done in the KBase environment (59). In summary, following quality trimming and adapter removal, the remaining 126 M paired-end reads were assembled with MetaSPAdes (v3.13.0) and a minimum contig size of 500 bp (60). The assembled contigs were binned using the MetaBAT2 (v1.7) binning algorithm (61). The quality of the resulting ten bins was evaluated using the CheckM (v1.0.18) software package (62). Taxonomy was assigned to individual bins using the Genome Taxonomy Database Toolkit (GTDB-Tk), which utilizes domain-specific, concatenated marker protein reference trees, relative evolutionary divergence and average nucleotide identity for taxonomic classification (63). Structural and functional annotation of binned contigs was achieved with NCBI’s prokaryotic annotation pipeline (64, 65). Additional annotations of unbinned contigs were obtained using the Prokka software package (66).

### Data availability

16S rRNA amplicon sequencing reads and shotgun metagenome reads used for assembly and binning of individual genomes have been deposited to NCBI’s sequencing reads archive and the annotated draft genomes have been deposited at GenBank. The data can be accessed under the bioproject PRJNA644413.

## ACKNOWLEDGEMENT

We thank Manuel Castelan, Benny Chacko and Yao Xiong for their thoughtful discussions and technical support throughout this project. We also greatly acknowledge the assistance provided by Kingsley Oparaodu in obtaining several of the oil field samples.

