## Supplementary material for "Severe corrosion of carbon steel in oil field produced water can be linked to methanogenic archaea containing a special type of [NiFe] hydrogenase"

**Table S1** Top 25 hits for a protein sequence BLAST (BlastP) search of MicH against NCBI's non-redundant protein database.

| Putative function | Closest taxonomic classification | Total BlastP score | Query coverage (%) | E-value | Sequence identity (%) | Accession No. |
|---|---|---|---|---|---|---|
| nickel-dependent hydrogenase large subunit | *Methanococcus maripaludis* OS7 | 1026 | 100 | 0.0E+00 | 100.0 | WP_119846095.1 |
| hypothetical protein | *Methanobacterium* sp. | 946 | 98 | 0.0E+00 | 93.5 | RJS49388.1 |
| hypothetical protein | *Methanobacteriales* | 887 | 98 | 0.0E+00 | 90.7 | PKL67834.1 |
| nickel-dependent hydrogenase large subunit | *Methanobacterium congolense Buetzberg* | 884 | 98 | 0.0E+00 | 90.7 | WP_071906935.1 |
| hypothetical protein | *Methanopyrus* sp. SNP6 | 340 | 96 | 9.0E-108 | 40.7 | WP_148689234.1 |
| hypothetical protein | *Bacteria* | 176 | 96 | 2.0E-45 | 28.5 | NLI99272.1 |
| Ni/Fe hydrogenase subunit alpha | Ca. *Korarchaeota* | 170 | 96 | 2.0E-43 | 28.8 | HDD68771.1 |
| Fe hydrogenase | *Phycisphaerae* | 170 | 98 | 4.0E-43 | 29.0 | KPK74792.1 |
| Ni/Fe hydrogenase subunit alpha | *Desulfosporosinus* sp. Sb-LF | 168 | 96 | 2.0E-42 | 28.0 | WP_135380469.1 |
| Ni/Fe hydrogenase subunit alpha | *Nitrosomonadales* | 168 | 96 | 2.0E-42 | 29.5 | PWB56486.1 |
| Ni/Fe hydrogenase subunit alpha | *Sulfuritalea hydrogenivorans* | 167 | 98 | 2.0E-42 | 29.4 | WP_041099025.1 |
| Ni/Fe hydrogenase subunit alpha | *Archaea* | 167 | 96 | 4.0E-42 | 28.6 | HEA10786.1 |
| Ni/Fe hydrogenase subunit alpha | *Nitrosospira* sp. Nsp13 | 166 | 96 | 7.0E-42 | 29.2 | WP_090907184.1 |
| Ni/Fe hydrogenase subunit alpha | *Nitrosospira multiformis* | 166 | 96 | 8.0E-42 | 28.7 | WP_107761584.1 |
| Ni/Fe hydrogenase subunit alpha | candidate division WOR-3 | 165 | 97 | 1.0E-41 | 28.2 | HHR48257.1 |
| Ni/Fe hydrogenase subunit alpha | candidate division WOR-3 | 165 | 97 | 2.0E-41 | 27.9 | HGU47450.1 |
| NADP oxidoreductase | *Nitrosomonadales* | 165 | 96 | 2.0E-41 | 29.0 | ODT84812.1 |
| Ni/Fe hydrogenase subunit alpha | *Nitrosomonadales* | 165 | 96 | 2.0E-41 | 29.5 | TFH12556.1 |
| Ni/Fe hydrogenase subunit alpha | candidate division WOR-3 | 164 | 97 | 4.0E-41 | 28.7 | HGK62977.1 |
| Ni/Fe hydrogenase subunit alpha | Ca. *Bathyarchaeota* | 163 | 96 | 6.0E-41 | 28.5 | HGD66108.1 |
| Ni/Fe hydrogenase subunit alpha | *Nitrosospira lacus* | 164 | 96 | 8.0E-41 | 29.9 | WP_004174406.1 |
| nickel-dependent hydrogenase large subunit | *Aquisphaera* sp. JC650 | 163 | 98 | 9.0E-41 | 29.4 | WP_152053939.1 |
| F420-non-reducing hydrogenase subunit A | *Lokiarchaeum* sp. GC14_75 | 163 | 96 | 9.0E-41 | 27.9 | KKK40681.1 |
| Ni/Fe hydrogenase subunit alpha | *Firmicutes* | 163 | 98 | 9.0E-41 | 27.9 | PKM81450.1 |
| Ni/Fe hydrogenase subunit alpha | *Anaerolineae* | 163 | 96 | 1.0E-40 | 30.3 | HGY48290.1 |

**Table S2** Microorganisms used as negative controls to evaluate *micH* qPCR assay.

| Organism name | Genome Accession No. | Origin | Country | Genome status | *micH* detected |
|---|---|---|---|---|---|
| *Archaeoglobus fulgidus* ATCC 49558 | NC_000917.1 | Hot Spring | Italy | Complete | − |
| *Burkholderia thailandensis* ATCC 700388 | NC_007651.1 | Rice field soil | Thailand | Complete | − |
| *Desulfovibrio vulgaris* ATCC 29579 | NC_002937.3 | Wealden clay | United Kingdom | Complete | − |
| *Eubacterium limosum* ATCC 8486 | NZ_CP019962.1 | Human intestinal content | Germany | Complete | − |
| *Geobacter metallireducens* ATCC 53774 | NC_007517.1 | Fresh water sediment, Maryland | United States | Complete | − |
| *Geobacter sulfurreducens* ATCC 51573 | NC_002939.5 | Surface sediment, Oklahoma | United States | Complete | − |
| *Methanobacterium bryantii* ATCC 33272 | NZ_LMVM00000000.1 | Syntrophic culture of "*Methanobacterium omelianskii*" | NA | Draft | − |
| *Methanocaldococcus jannaschii* ATCC 43067 | NC_000909.1 | Submarine hydrothermal vent | Pacific Ocean, East Pacific Rise | Complete | − |
| *Methanococcus voltae* ATCC BAA-1334 | CP002057.1 | Salt marsh, Florida | United States | Complete | − |
| *Methanoculleus bourgensis* ATCC 43281 | NC_018227.2 | Tannery sewage sludge | France | Complete | − |
| *Methanomicrobium mobile* ATCC 35094 | NZ_JOMF00000000.1 | Bovine rumen | United States | Draft | − |
| *Nitrosomonas europaea* ATCC 19718 | NC_004757.1 | NA | NA | Complete | − |
| *Roseobacter denitrificans* ATCC 33942 | NC_008209.1 | Seaweed | NA | Complete | − |
| *Shewanella denitrificans* ATCC BAA-1090 | NC_007954.1 | Water column, 130 m depth | Baltic Sea | Complete | − |
| *Thiobacillus denitrificans* ATCC 25259 | NC_007404.1 | Soil, Texas | United States | Complete | − |

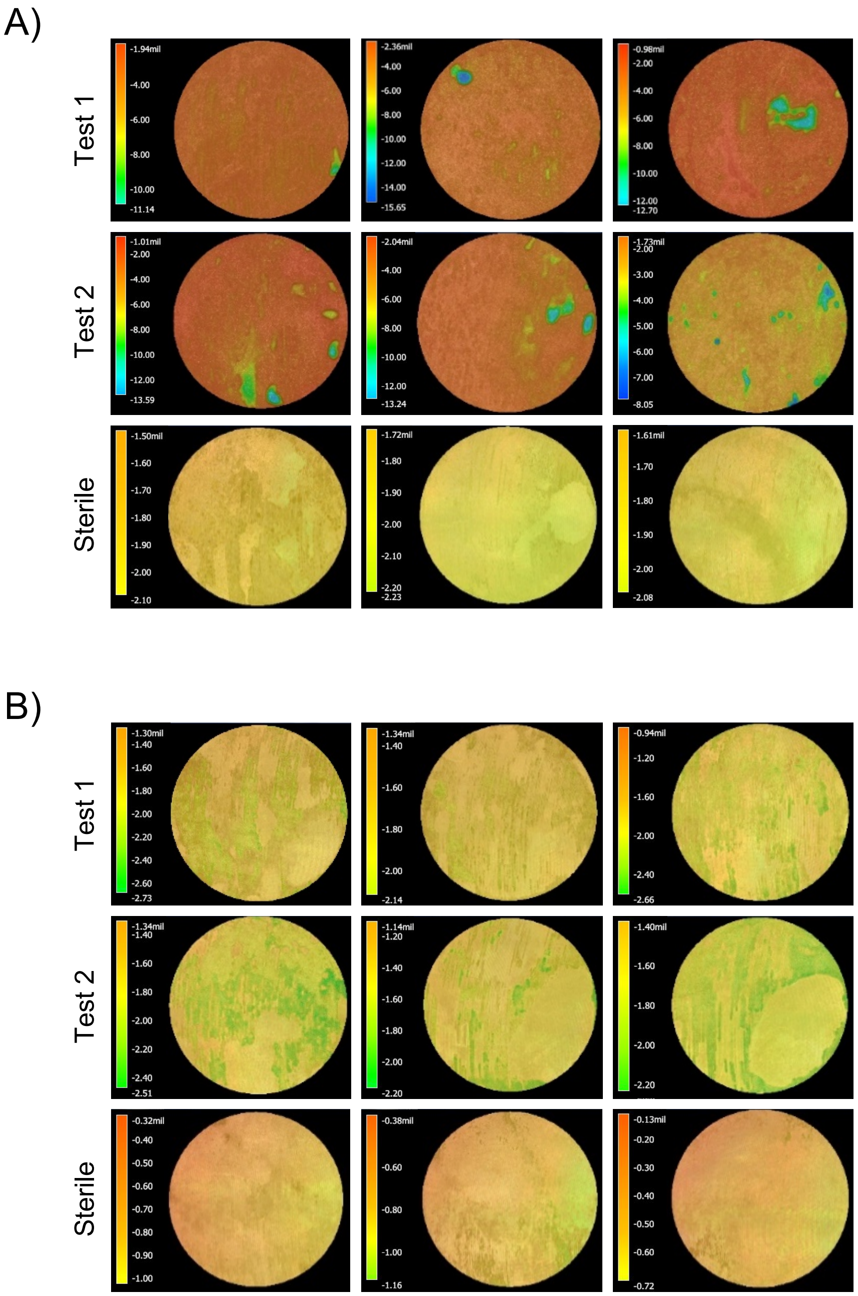


**Figure S1** Surface topography of cleaned X52 carbon steel weight loss corrosion coupons after exposure to produced water at A) 32°C and B) 60°C. See Figure 2A for corresponding weight loss corrosion data.

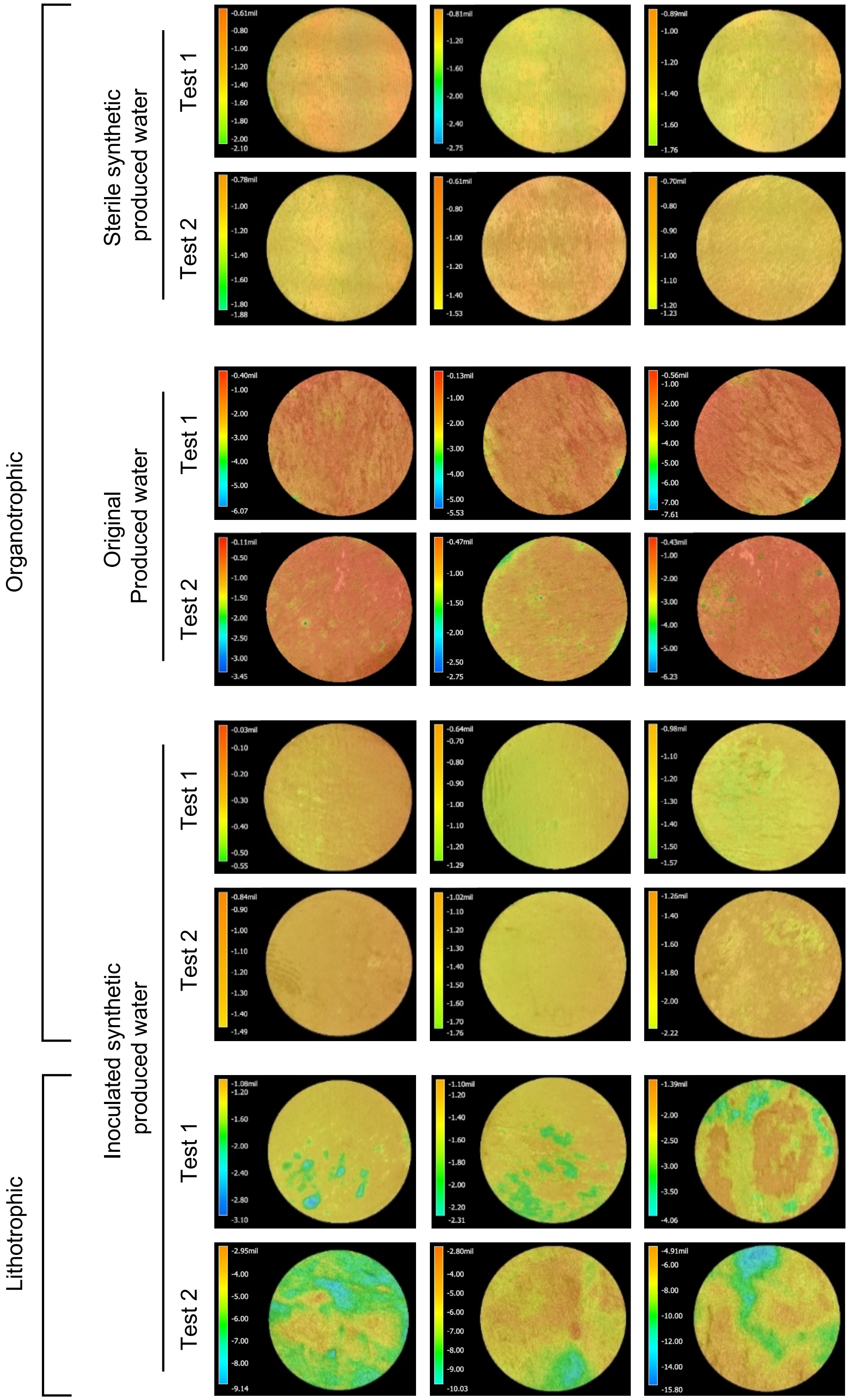


**Figure S2** Surface topography of cleaned X52 carbon steel weight loss corrosion coupons after exposure to original produced water, sterile or inoculated synthetic produced water at 32°C under organotrophic or lithotrophic conditions. See Figure 2D for corresponding weight loss corrosion data.

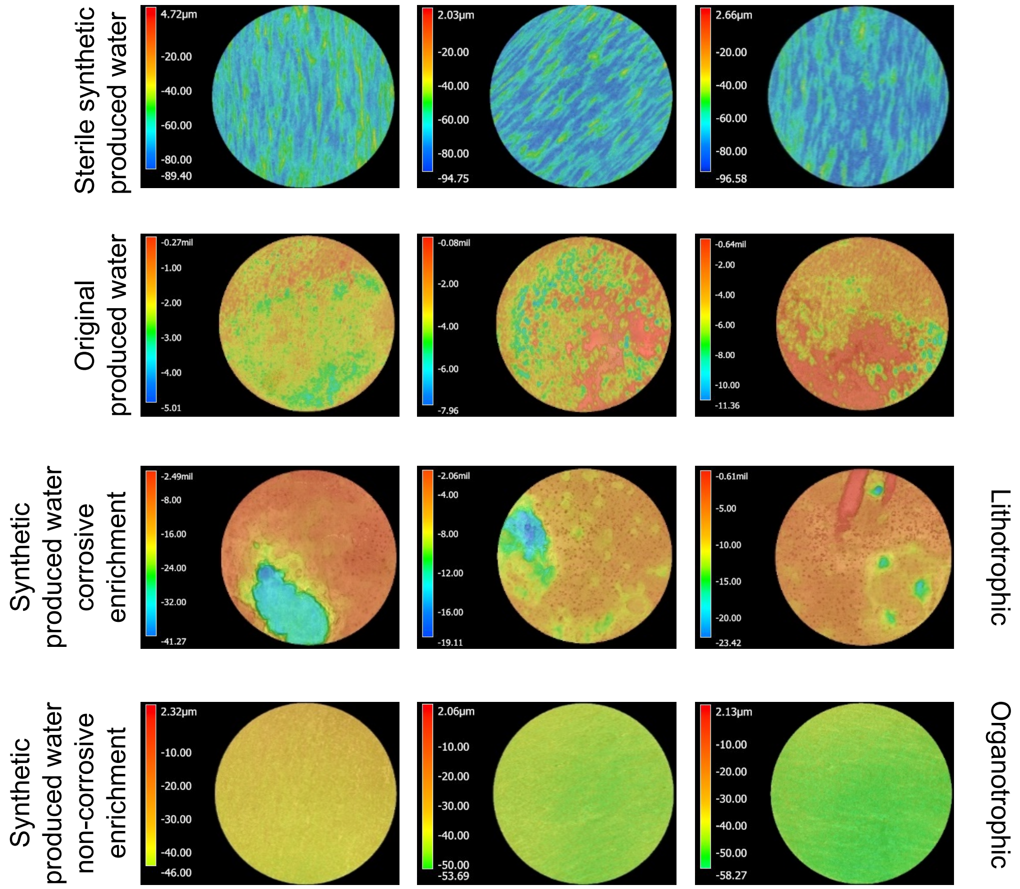


**Figure S3** Surface topography of cleaned X52 carbon steel weight loss corrosion coupons after exposure to produced water or synthetic produced water under simulated pipeline conditions in customized reactors. See Figure 3D for corresponding weight loss corrosion data.

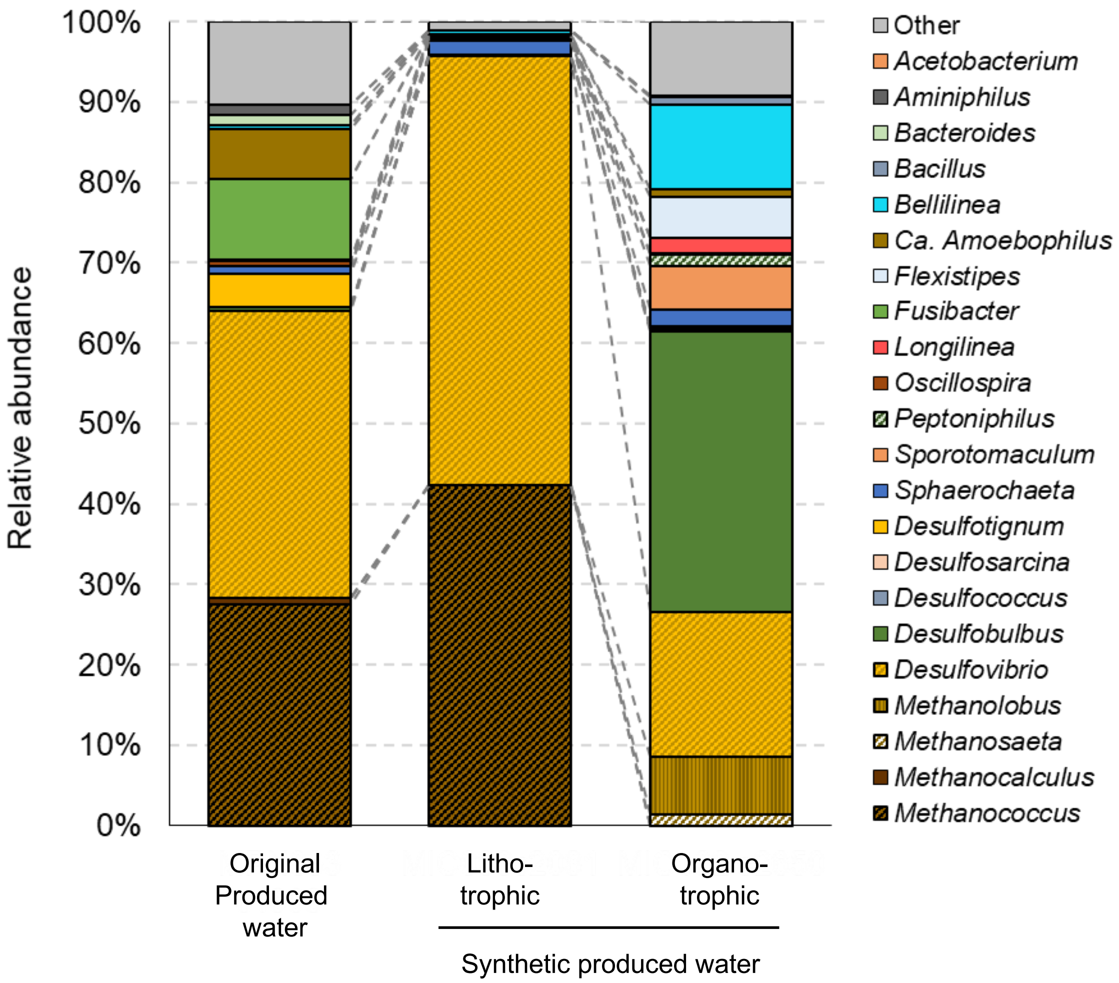


**Figure S4** Microbial community composition on carbon steel coupon surfaces after incubation in customized reactors under simulated pipeline conditions. See Figure 3D for weight loss corrosion rates.

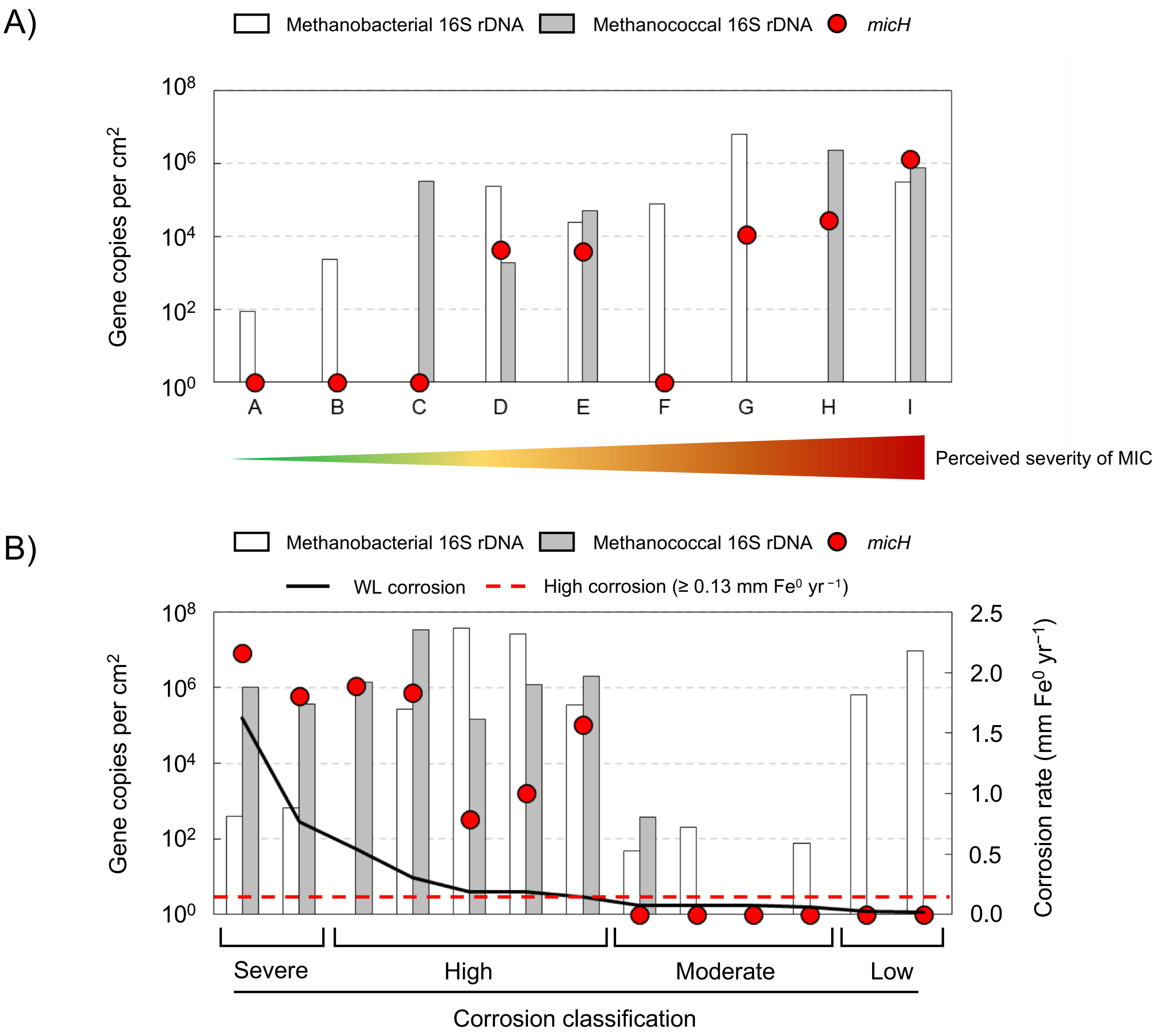


**Figure S5** Comparison of corrosion rate, *micH* gene copies and estimated methanococcal and methanobacterial 16S rRNA gene copies in A) pigging debris samples shown in Figure 1 and B) laboratory tests shown in Figures 2 and 3. Genus specific estimated 16S rRNA gene copies were calculated from total 16S rRNA gene copy numbers obtained by qPCR and relative community composition data from 16S rRNA gene amplicon sequencing.

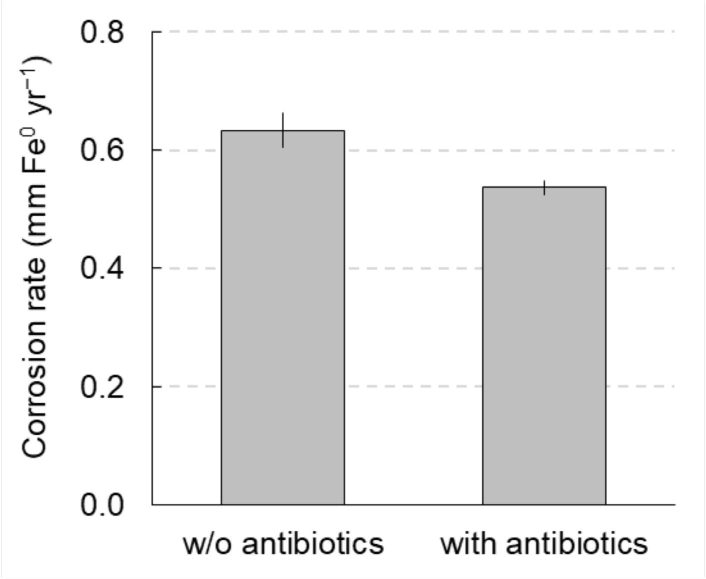


**Figure S6** Comparison of corrosion rate in kettle tests using synthetic produced water treated with or without (w/o) antibiotics.

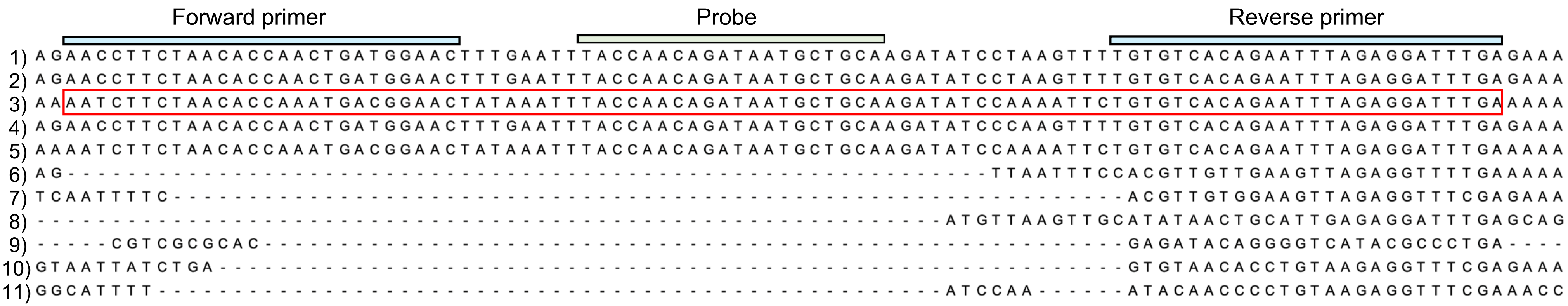


**Figure S7** Alignment of *micH* gene region used for primer and probe design (sequences 1 – 5). Primer and probe binding sites are shaded with different colors and the synthetic fragment used for quantification is edged in red. Alignment of large subunits of remotely related hydrogenases found in methanogens not carrying the *micH* gene (sequence 6 – 11) showed the desired specificity of the selected primers and probe. Sequence 1) *micH* (H0S71_01960) *Methanococcus maripaludis* MIC098Bin5 metagenome (JACCQJ000000000); 2) *micH* (MMOS7_11600) *Methanococcus maripaludis* OS7 (AP011528.1); 3) *micH* (CVV28_03670) *Methanobacteriales* groundwater metagenome (PGYO01000002.1); 4) *micH* (CIT03_03910) *Methanobacterium* bioreactor metagenome (NPZC01000005.1); 5) *micH* (MCBB_1253) *Methanobacterium congolense* Buetzberg (LT607756.1); 6) MMP_RS04285 *Methanococcus maripaludis* S2 (NC_005791.1); 7) MVOL_RS02900 *Methanococcus voltae* A3 (NC_014222.1); 8) MJ_RS06365 *Methanocaldococcus jannaschii* DSM 2661 (NC_000909.1; 9) MBBA_RS09640 *Methanoculleus bourgensis* (NZ_LT549891.1); 10) MVOL_RS02915 *Methanococcus voltae* A3 (NC_014222.1); 11) MMP_RS07120 *Methanococcus maripaludis* S2 (NC_005791.1). Sequences were aligned with MUSCLE in MEGA7 (Reference: Kumar S, Stecher G, Tamura K (2016) MEGA7: Molecular Evolutionary Genetics Analysis Version 7.0 for BIGGER Datasets. Mol Biol Evol 33: 1870–1874. doi:10.1093/molbev/msw054).

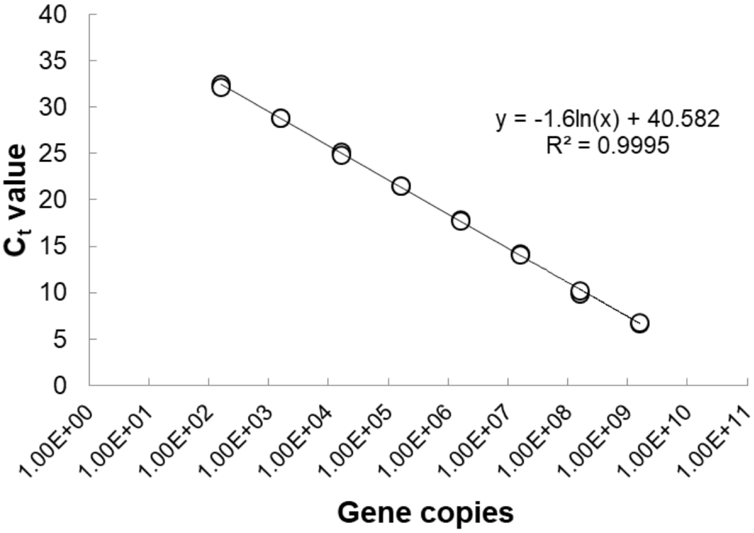


**Figure S8** Calibration curve of quantitative PCR using a synthetic fragments of *micH* gene as target and the newly developed primer pair and Taqman™ probe.
